# Pax7 pioneer factor action requires both paired and homeo DNA binding domains

**DOI:** 10.1101/2020.10.09.332015

**Authors:** Audrey Pelletier, Alexandre Mayran, Arthur Gouhier, James G Omichinski, Aurelio Balsalobre, Jacques Drouin

## Abstract

The pioneer transcription factor Pax7 contains two DNA binding domains (DBD), a paired and a homeo domain. Previous work on Pax7 and the related Pax3 had shown that each DBD can bind a cognate DNA sequence, thus defining two targets of binding and possibly modalities of action. Genomic targets of Pax7 pioneer action leading to chromatin opening are enriched for composite DNA target sites containing juxtaposed binding sites for both paired and homeo domains. The present work investigated the implication of both DBDs in pioneer action. We now show that the composite sequence is a higher affinity Pax7 binding site compared to either paired or homeo binding sites and that efficient binding to this site involves both DBDs. We also show that a Pax7 monomer binds composite sites and that methylation of cytosines within the binding site does not affect binding, which is consistent with pioneer action exerted at methylated DNA sites within nucleosomal heterochromatin. Finally, introduction of single amino acid mutations in either the paired or homeo domain that impair binding to cognate DNA sequences showed that both DBDs must be intact for pioneer action. In contrast, only the paired domain is required for low affinity binding of heterochromatin sites. Thus, Pax7 pioneer action on heterochromatin requires unique protein:DNA interactions that are more complex compared to its simpler DNA binding modalities at accessible enhancer target sites.

**Significance Statement:** Pioneer transcription factors have the unique ability to recognize DNA target sites within closed heterochromatin and to trigger chromatin opening. Only a fraction of the heterochromatin recruitment sites of pioneers are subject to chromatin opening. The molecular basis for this selectivity is unknown and the present work addressed the importance of DNA sequence affinity for selection of sites to open. The pioneering ability of the pioneer factor Pax7 is not strictly determined by affinity or DNA sequence of binding sites, nor by number or methylation status of DNA sites. Mutation analyses showed that recruitment to heterochromatin is primarily dependent on the Pax7 paired domain whereas the ability to open chromatin requires both paired and homeo DNA binding domains.

---

Pioneer factors are a subset of transcription factors (TFs) that have been dubbed with the unique property of being able to “open chromatin structure” (1-3). While pioneers have the typical TF domain organisation found in most TFs including DNA binding and transactivation domains, the molecular basis of their ability to “open chromatin” remains largely elusive. In fact, recent work clearly separated the chromatin opening function from the pioneer ability and showed that the former can be conferred through recruitment of a nonpioneer factor: this was shown for the pioneer Pax7 that is required to implement the melanotrope enhancer repertoire (4, 5) but that is dependent on the cell determination TF Tpit for chromatin opening (6). These findings thus limit the unique pioneer property to recognition of DNA binding sites within heterochromatin. A large body of work also showed that pioneer TFs can bind their target DNA sequence within nucleosomes that would presumably be tightly packed within heterochromatin (7, 8). Whereas the ability to recognize target DNA within nucleosomal structures appears to be a prerequisite for pioneer action, this in itself is not sufficient to explain the specificity of pioneer action nor to explain how initial sequence recognition is linked to chromatin remodelling.

Various pioneers activate novel enhancer repertoires that control cell specific gene expression programs. This was shown for the pioneers FoxA (9, 10), Pax7 (4, 5), C/EBPα (11), EBF1 (12, 13), Ascl1 (14), NeuroD1 (15), and GATA3 (16). For all these factors, with the exception of Pax7, the DNA binding sites for the TF at pioneered enhancers (hereafter, we refer to ‘pioneered enhancers/sites’ as those that require chromatin opening for activity) appeared similar to their previously defined transcriptional target sites. In contrast, the initial work on Pax7 pioneer target sites revealed enrichment for a unique composite target sequence that includes motifs recognized by each DNA binding domain (DBD) present in Pax7 (5). Indeed, Pax7 contains two DBDs, a paired domain (PD) and a homeo (HD) domain, and each recognizes different prototypical DNA target sites. Prior work on the DNA binding properties of Pax7 and the closely related Pax3 used cognate binding sites for either the PD or the HD (17), and these appear to show independent binding by the PD or HD domains. However, close interaction of the two DBDs appears important for optimal DNA binding (18, 19). Indeed, DNA binding by either the PD or HD causes a structural change in the other DBD (20). In this context, recognition of a unique composite DNA sequence containing binding motifs for both PD and HD enriched at pioneered targets suggest that Pax7 pioneer action may involve a unique conformation or binding requirement.

It is presently unclear what DNA recognition features may be critical for pioneer target recognition and action. Although higher affinity for DNA sites present at pioneer targets may be a mechanism to trigger structural changes at nucleosomal sites, various studies have shown the ability of pioneer TFs to recognize subsets of their target sequences by recognition of partial and presumably lower affinity DNA motifs. This was proposed to represent a “scanning” ability of pioneers to probe putative DNA targets within nucleosomal heterochromatin (21). This feature results in large numbers of pioneer-bound sites that are unproductive for chromatin remodelling and were thus labelled as “resistant” (4) or as unchanged (21, 22). The contribution of DNA binding affinity for pioneer action thus remains unclear. It was recently shown that the FoxA pioneer factor requires a core histone interaction domain for chromatin opening (23). However, the scanning ability and core histone interaction do not explain the selection of site subsets that are productive for chromatin opening.

The present work investigated the DNA binding properties of Pax7 in relation to its various target DNA sequences both *in vitro* using gel mobility shift and *in vivo* using ChIPseq. The highest *in vitro* binding affinity was observed with a composite Pax7 target that includes DNA motifs for recognition by both PD and HD and this composite sequence is enriched at pioneered enhancers. The composite DNA site is bound *in vitro* by a Pax7 monomer and the binding affinity is unchanged when the composite sequence contains a methylated CpG, a feature that would be expected of pioneers targeting regulatory sequences within nucleosomal heterochromatin. Finally using single amino acid substitutions of key residues required for DNA binding of either PD or HD, we show that the DNA binding activity of both domains are required for pioneer action whereas the PD alone is sufficient for binding to low affinity resistant sites. In summary and in contrast with the ability of each DNA binding domain to bind its target sequence in naked DNA, the dependence on both domains in order to trigger chromatin opening at pioneered sites suggests that a unique conformation of the Pax7 DBDs is needed for the initial events leading to chromatin opening.

## RESULTS

### Strongest in vitro Pax7-binding motif is enriched at pioneered target sites

Prior studies (reviewed in (17)) on the DNA binding properties of the related Pax TFs Pax3 and Pax7 (Fig.1A) revealed cognate binding sites for either of their two DBDs, the PD or the HD domain (Fig.1B). Genome-wide assessment of these TF binding sites obtained by ChIPseq showed enrichment of the PD target sequence core motif TGAC and of different associations of the TAAT motif recognized by the HD (5, 24). We also reported that Pax7 pioneered target enhancers show enrichment of the composite motif that contains juxtaposed PD and HD motifs (5). Extensive characterization of the Pax7 ChIPseq peaks defined different genomic recruitment sites (4). These included binding at putative enhancer sites that are already accessible (Constitutive sites), at sites where Pax7 binding results in transcriptional activation (Activated sites revealed by p300 recruitment), at Resistant sites where Pax7 is recruited but without any changes in chromatin organisation, and finally at pioneered sites that either become fully transcriptionally competent (Pioneer, p300-positive) or Primed (p300-negative). Pax7 motif frequencies for these different subcategories show that the fully activated Pioneer sites have the highest frequency for all motifs (Fig.1C, D) and this correlates with this category of peaks exhibiting the strongest Pax7 ChIPseq signals of all categories (4) as supported by the highest median p values for this subset (Fig. 1C). The Pioneer subset also has the highest combined frequencies of all different Pax7 motifs (Fig.1E) consistent with the frequent occurrence of more than one motif under each peak (Fig.1F).

**Fig. 1.**
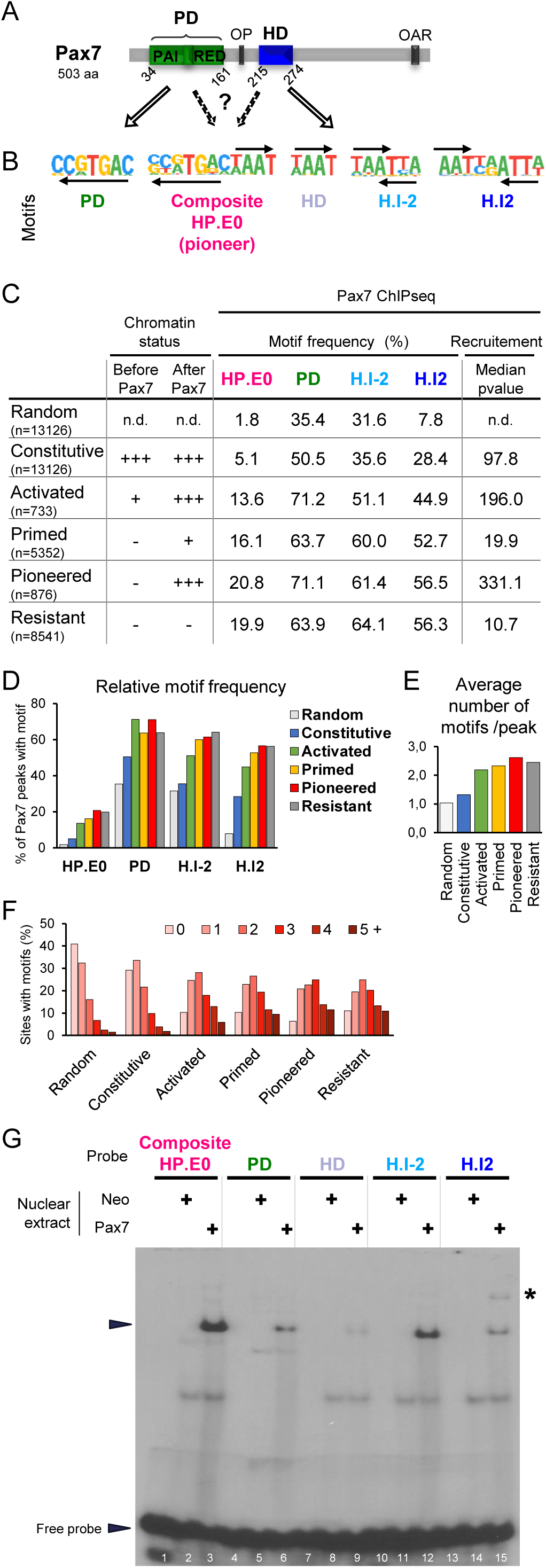
Contribution of paired (PD) and homeodomain (HD) DNA binding domains to Pax7 binding. (A) Schematic representation of mouse Pax7 structure (Uniprot P47239-1) including the PD and HD DNA binding domains. The positions of two other conserved domains is also indicated: OP, a conserved octapeptide present in the linker region between the two DNA binding domains and OAR, a conserved tetradecapeptide initially identified in the transcription factors otp, aristaless and rax. (B) Consensus recognition motifs identified in Pax7 ChIPseq (5) and related to recognition by either PD or HD as suggested by arrows linking panels A and B. The composite HP.E0 motif contains everted sequences of the PD and HD recognition motifs and this composite was identified as enriched under Pax7 peaks that trigger pioneer chromatin opening (5). (C) Distribution of Pax7 interaction motifs in subsets of Pax7 ChIPseq peaks defined by their changes (or not) in pituitary AtT20 cells before/after Pax7 expression. These categories were defined and discussed extensively in (4), and the subsets presented here (n indicated between parentheses) were trimmed to only include peaks with stringently defined properties compared to other subsets. For reference, the chromatin accessibility (ATAC) status of these subsets is indicated. The average Pax7 recruitment strength for each subset is indicated by the median P value of peaks present in each subset. (D) Relative frequencies (% of peaks containing indicated motifs) for each subset of peaks. (E) Average total number of all motifs per peak. (F) Distribution of number of motifs per peak for each subset of Pax7 ChIPseq peaks. (G) Binding of Pax7 to its different DNA targets. Electrophoretic Mobility-Shift Assay (EMSA) was used to assess the in vitro binding of Pax7 stably expressed in AtT20 cells (4). Similar DNA probes (probe sequences in Fig. S1A) were incubated with either control AtT20-neo or AtT20-Pax7 cell nuclear extracts for analyses by EMSA. A single specific band is observed (arrowhead) for each probe but with widely varying intensities. Only the H.I2 palindromic probe shows an additional band (*) that is likely constituted of Pax7 homodimers.

In order to compare Pax7 binding to each target type motifs, we did electrophoretic mobility shift assays (EMSA) using DNA probes corresponding to each motif (Fig. S1A) and nuclear extracts from AtT20 cells expressing a Flag-tagged Pax7 or an empty vector (Neo). Complexes present in EMSA using Pax7 compared to Neo cells revealed a single complex present with all probes (arrowhead in Fig. 1G). The apparent strength/stability of this Pax7 complex is however very different for different probes: the highest apparent affinity site is the composite HP.E0 site followed, in decreasing order, by the inverted TAATTA palindrome H.I-2, the PD motif, the TAAT palindrome H.I2, and the single HD. The only other Pax7-specific complex is observed with the H.I2 probe (asterisk in Fig. 1G). This slower migrating band is a likely Pax7 dimer as this sequence was previously shown to form dimers with both Pax3 and Pax7 (25). We did competitive binding experiments in order to assess the relative affinities of complexes formed with the different probes and this supported the interpretation of the highest affinity for the composite followed by PD and HD sites (Fig.S1B). The presence of Pax7 in the purported Pax7-DNA complexes was ascertained by supershift EMSA experiments using the Flag antibody: both monomer and dimer Pax7:DNA complexes were indeed confirmed to contain the Flag-Pax7 chimeric protein (Fig. S1C).

### Prevalent role of Pax7 paired (PD) domain for DNA binding

In order to assess the relative roles of PD and HD domains for DNA binding, we produced and purified polypeptides containing different Pax7 domains from over-expressing *E. Coli*. Whereas the PD domain (residues 35-158 of mouse Pax7) is sufficient for in vitro DNA binding to the composite HP.E0 and PD probes, the HD domain (aa 215-273) does not bind significantly (Fig. 2A, lanes 1, 3, 5, 7). Addition of the linker sequence present between PD and HD to the PD polypeptide appeared to stabilize PD probe binding (lane 6 compared to 5) but the strongest binding is observed for a polypeptide containing both PD and HD (PD.HD, aa 35-273) together with intervening linker sequences (Fig. 2A, lanes 4 and 8). On its own, the HD polypeptide exhibited strong binding only to the palindromic H.I2 probe compared to the core HD or to the HD H.I-2 probes (Fig. 2A, lanes 19 compared to 11, 15). The PD:HD polypeptide showed the strongest binding to all probes although this is marginal for the simple HD motif (lanes 4, 8, 12, 16, 20). Further, this PD:HD polypeptide presents an additional band with the H.I2 palindromic probe (lane 20) consistent with homodimer binding reported for both Pax3 and Pax7 with this sequence (25). Collectively, these data indicate that the PD domain is primarily responsible for high affinity DNA binding and that the HD domain may contribute stabilization and specificity of binding, in particular with the composite HP.E0 probe. On the other hand, the HD may increase the repertoire of Pax7 binding sites to include inverted palindromic target sequences of the H.I2 type.

**Fig. 2.**
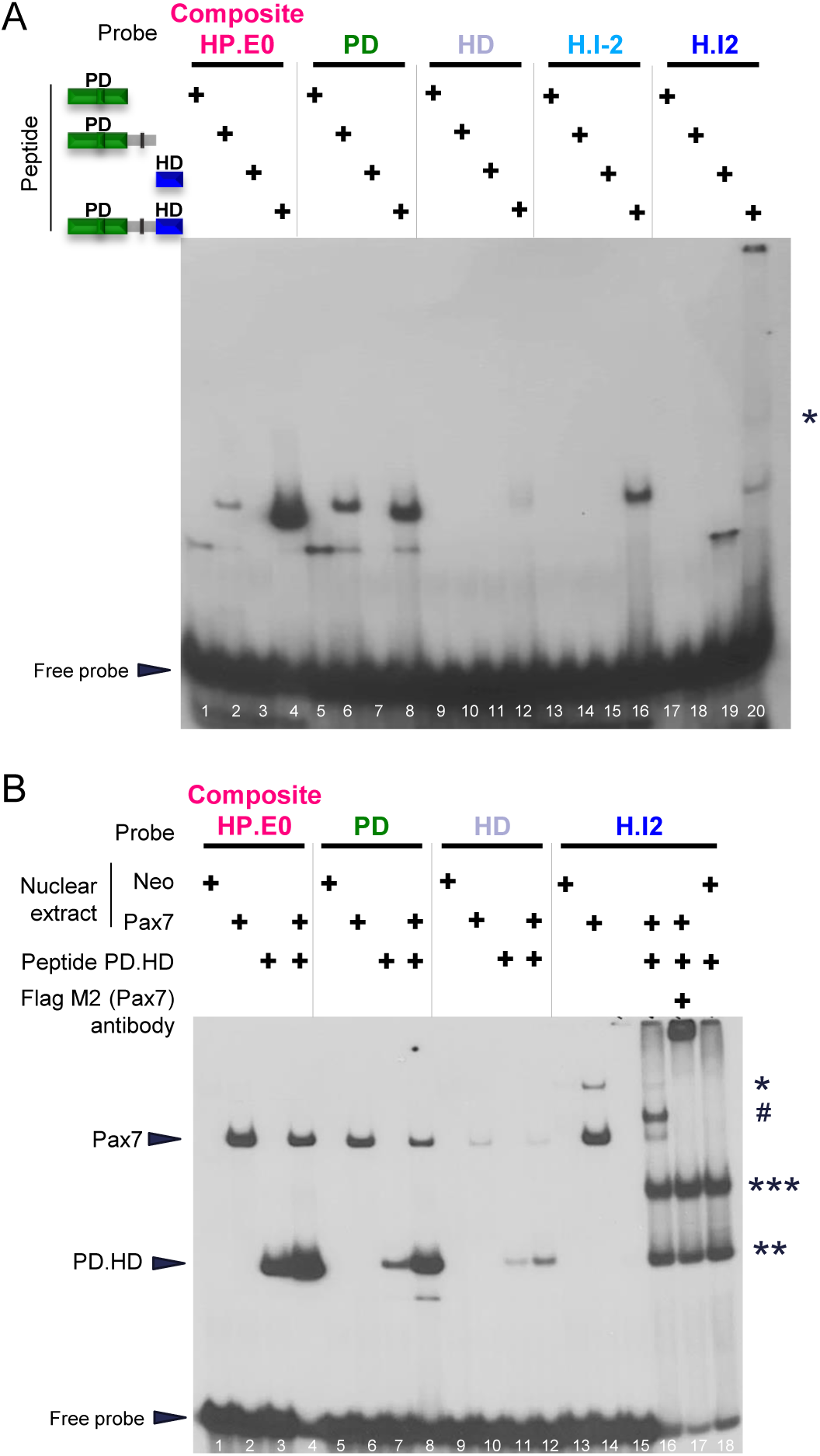
Relative roles of Pax7 PD and HD domains in DNA binding. (A)In vitro binding (EMSA) of various Pax7 polypeptides produced in E. coli to contain the represented portions of the Pax7 DNA binding domains. The purified polypeptides were incubated with ^32^P-labelled probes before electrophoresis. For each probe, the strongest band likely represents monomer Pax7 polypeptide binding except for the H.I2 probe where a likely dimer band is observed (*). (B)EMSA mixing experiment showing that Pax7 monomers bind the composite site. EMSA was performed for each DNA probe using either AtT20 nuclear extracts expressing (or not, neo) Pax7 with / without the PD.HD polypeptide as indicated. Arrowheads indicate the positions of wild-type Pax7 and PD.HD polypeptide monomer complexes. For the H.I2 probe, dimer bands are also observed for Pax7 (*) and PD.HD (***) together with appearance of a new band (#) in the combined reaction (lane 16) that likely represents a heterodimer between intact Pax7 and PD.HD polypeptide. The presence of Flag-Pax7 in complexes is ascertained by supershifts with the Flag M2 antibody.

### Pax7 binds the composite site as monomer

Binding of Pax7 to the composite element may involve the two DBDs of the same Pax7 molecule or the PD of one associated as dimer with the HD of another Pax7 molecule. In order to address this question, we performed EMSA using AtT20-Pax7 nuclear extracts with and without addition of the bacterially-produced PD.HD polypeptide. These mixing experiments (Fig. 2B) revealed independent binding of the PD.HD polypeptide and AtT20-expressed Pax7 to the composite (lanes 2-4), PD (lanes 6-7) and HD (lanes 11-13) probes. In contrast, binding of the full-length Pax7 to the palindromic H.I2 probe revealed two bands deemed to represent monomers and dimers (Fig. 2B, lane 14). The PD.HD polypeptide also formed two complexes with the slower migrating band (labelled ***) deemed to represent a dimeric complex that is not observed with the other probes (Fig. 2B, lane 18). When both PD.HD polypeptide and full-length Pax7 are present together, a new band (labelled #) was observed and interpreted as consisting of one molecule of full-length Pax7 together with one molecule of the PD.HD (Fig. 2B, lane 16). This new complex together with the Pax7-containing complexes were all supershifted by addition of the anti-Flag antibody confirming presence of the AtT20-expressed Pax7 in these complexes (Fig. 2B, lane 17). Taken together, these experiments are consistent with the interpretation that a monomer of Pax7 is sufficient to bind the composite target sequence with the PD and HD of the same Pax7 molecule likely to participate in close interactions in order to bind their juxtaposed target DNA sequences.

### Cytosine methylation does not interfere with Pax7 DNA binding

Prior to Pax7 binding, the pioneered enhancers are within nucleosomal heterochromatin (4) as determined by MnaseSeq (Fig. S2). Upon Pax7 binding, nucleosomes are excluded from the Pax7 pioneered sites (Fig. S2) and this correlates with the loss of histone H3 (4). The Pax7 pioneered sites are also within heterochromatin with high levels of CpG DNA methylation and this includes CpG residues that are within composite sites in some cases (4). These enhancers acquire stable epigenetic memory through demethylation of cytosine residues within targeted enhancers (Fig. 3A, B). Indeed, the average level of CpG methylation drops from > 90% to ∽30% after Pax7-dependent chromatin opening at the pioneered enhancers (Fig. 3B). These data suggest that cytosine methylation of DNA target sequences does not interfere with Pax7 interaction, and we directly assessed this by producing DNA probes with cytosine methylation of the CpG dinucleotide present within the composite Pax7 binding site (Fig. 3C). Probes containing methylcytosine on the sense (S, lane 6), the antisense (AS, lane 9) or both strands (lane 12) were compared to an unmethylated probe for Pax7 binding and Pax7 was found to bind equally well to all probes (Fig. 3D). This was also found to be the case for Pax7 binding to the methylated PD probe which showed weaker, but nonetheless methylation-independent, binding (Fig. 3D, lanes 13-25).

**Fig. 3.**
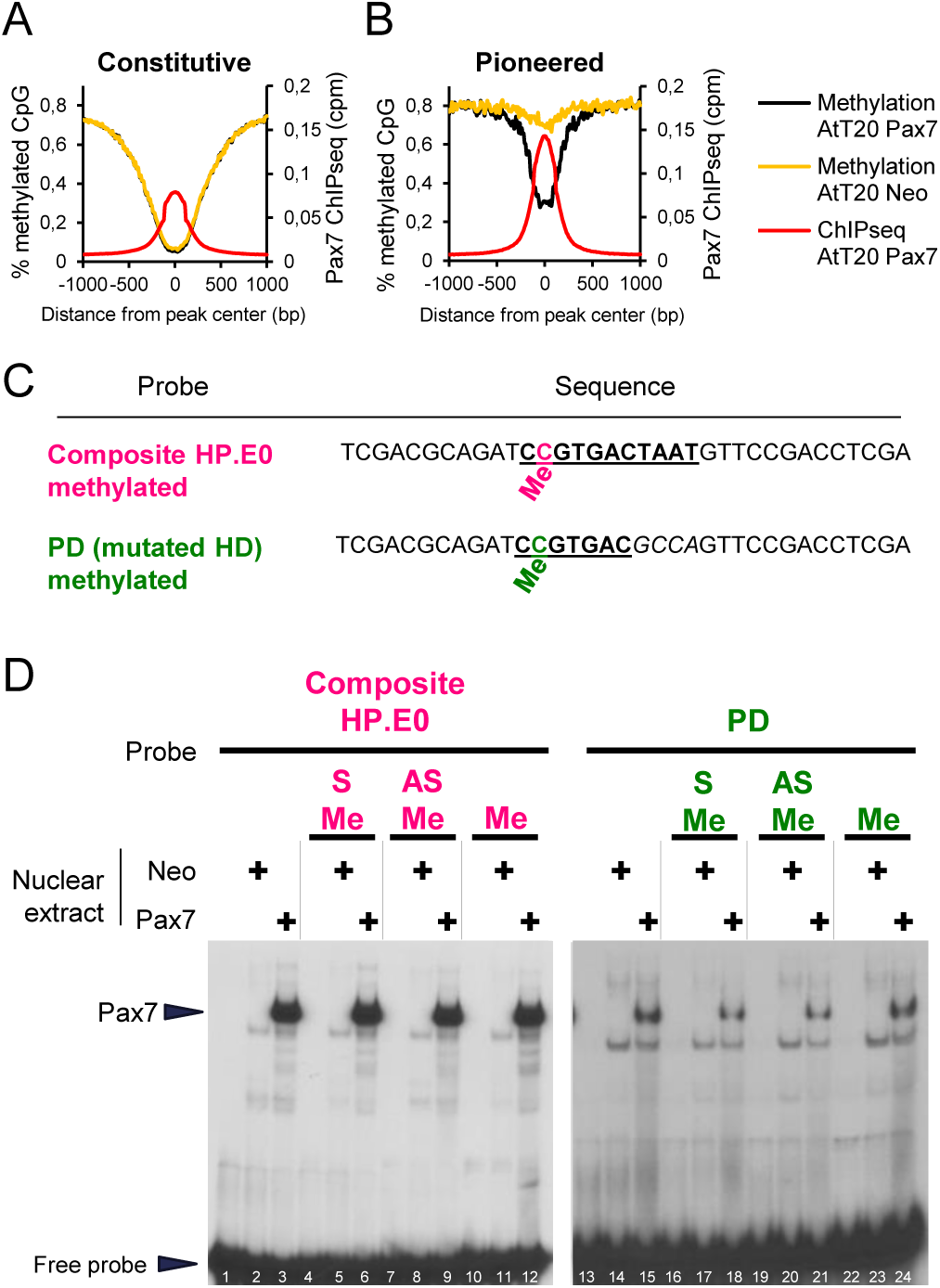
Pax7 binding is independent of DNA methylation. A, B) Average profiles of DNA methylation (% methylated CpG) around Pax7 ChIPseq peaks present at Constitutive (A) and Pioneered (B) enhancers. Data are shown for control AtT20-neo compared to AtT20-Pax7 cells and aligned with the average Pax7 ChIPseq profiles. (C)DNA sequences of probes used in EMSA to test the effect of DNA methylation on Pax7 binding. Methylated cytosine (C) residues are shown on sense (S) strand. (D)In vitro binding (EMSA) of Pax7 to either composite or PD probes methylated or not on sense (S), antisense (AS) or both (Me) strands.

The ability of Pax7 to bind its composite and PD sequences independently of cytosine methylation status is entirely consistent with the idea that this pioneer factor can recognize its target sequence within “closed” nucleosomal heterochromatin that is heavily methylated on cytosines.

### Both Pax7 DNA binding domains are required for pioneer activity

To ascertain the role of the two Pax7 DBDs in transcription and pioneer action, we introduced single amino acid mutations in either PD or HD (Fig. 4A). For the PD domain, the amino acids substitutions R56A and G43A were derived from Waardenburg syndrome mutations identified in the related PAX3 and shown to be loss-of-function mutations (26, 27), and for the HD domain, the critical recognition residue Serine 50 of the HD was changed to glutamic acid (S264E). These mutations were introduced in Flag-Pax7 and stably expressed in AtT20 cells as previously for Pax7 (5). The three Pax7 mutant proteins are expressed at similar levels (Fig.S3A). In vitro (EMSA), the R56L PD mutant did not bind the PD probe whereas the G43A PD mutant retained minimal binding activity (Fig. S3B, lanes 8 and 9). In contrast, the HD S264E mutant appeared to bind the PD probe slightly better than wild type Pax7 (lane 10 compared to 7). Strikingly, all three mutations had much less effect on binding to the composite probe with the only significant loss of binding activity observed for the PD R56L mutant (lanes 2-5). For HD probes H.I2 and H.I-2, the HD S264E mutant lost most binding activity but so did the PD mutants (lanes 16-25).

**Fig. 4.**
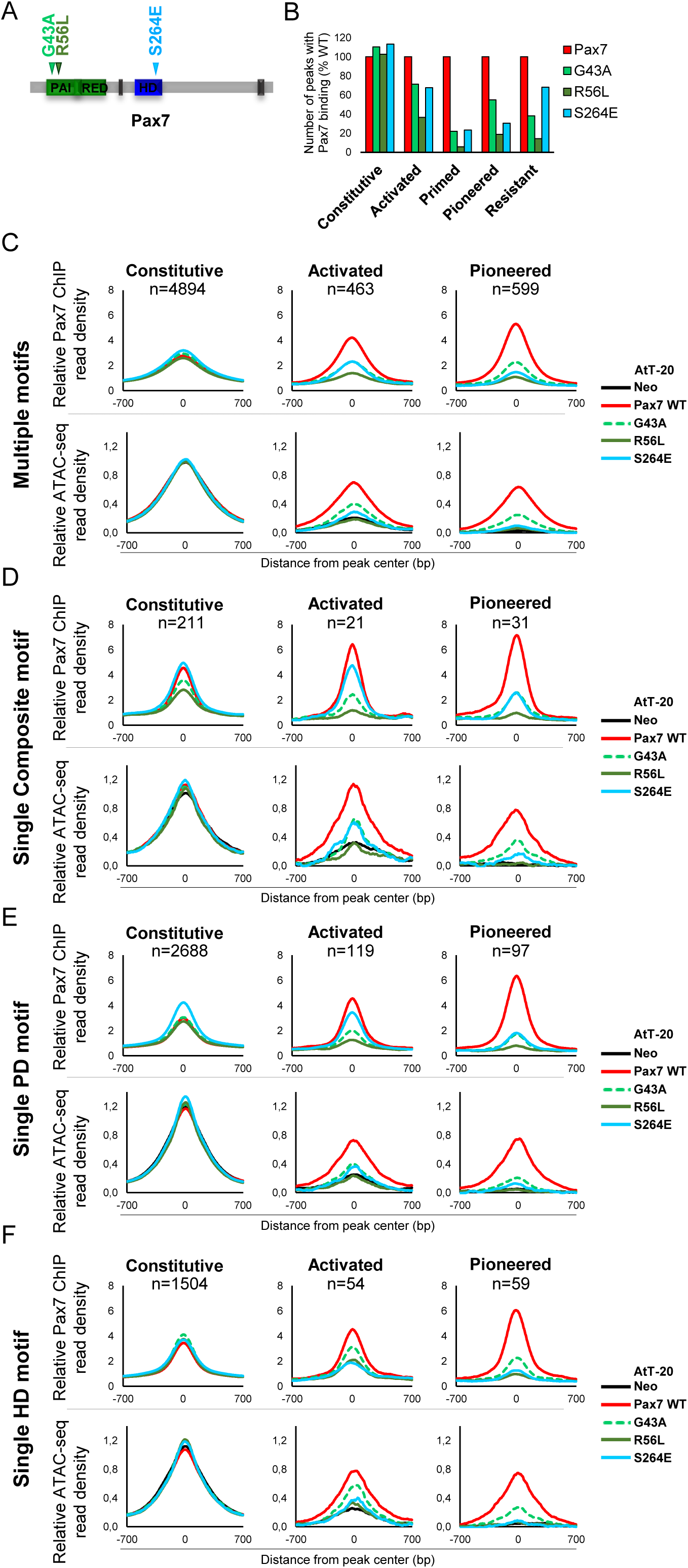
Intact and functional PD and HD domains are both required for recruitment and chromatin opening at Pax7 pioneer sites. (A)Schematic representation of Pax7 showing three amino acid substitutions assessed for recruitment by ChIPseq and chromatin opening by ATACseq. (B)Effect of three Pax7 mutations on ChIPseq recruitement at different subset of peaks as indicated. The number of peaks (% compared to wild-type Pax7) that retained significant binding are shown. (C)Average profiles of Pax7 ChIPseq and ATACseq signals for the indicated subsets of Pax7 peaks and for the subsets of peaks in each category that have multiple Pax7 recognition motifs. (D) Average profiles for the subsets of Pax7 peaks of the indicated categories that only contain one composite motif. (E)Average profiles for the subsets of Pax7 peaks of the indicated categories that only contain one PD motif. (F) Average profiles for the subsets of Pax7 peaks of the indicated categories that only contain one HD motif.

The transcriptional potential of these mutants was assessed by co-transfection into AtT-20 cells using a Luciferase reporter driven by six copies of the composite element or by the PC2 gene enhancer (5). For the hexamer reporter, transcriptional activity was only affected by the R56L mutation and not by the S264E HD mutation. These results (Fig. S3C) are consistent with the in vitro binding results (Fig. S3B). In contrast, the activity of the PC2 enhancer reporter was dependent on both PD and HD as evidenced by significant activity losses for all three mutants (Fig. S3C). The dependence on both the PD and HD for activity at this natural enhancer is thus more complex than the simpler in vitro binding patterns observed in EMSA.

The impact of the DBD mutations on *in vivo* genomic recruitment was assessed by ChIPseq (Fig.4B). It is quite striking to observe that key amino acid substitutions in either PD or HD failed to affect the recruitment of Pax7 to the Constitutive enhancers that are already in an open chromatin state: this likely reflects the strong contribution of protein-protein interactions for recruitment of Pax7 in the context of active enhancers. In contrast, recruitment to the pioneered enhancers is severely curtailed by all three mutations (loss of recruitment at ∽80% of sites for R56L, at ∽70% sites for S264E and ∽50% sites for G43A) with the G43A PD substitution having a partial effect compared to the R56L substitution (Figs. 4B and S3D). Thus, both DBDs must be intact for pioneer site binding whether considering the fully activated Pioneer sites or the Primed sites. Intermediate effects of the different mutations are observed at the group of so-called Activated enhancers, a group of enhancers with open chromatin where the co-activator p300 is recruited after Pax7 binding. Collectively, these data indicate that the G43A mutant is a partial loss-of-function whereas the other mutants appear complete nulls. Finally, it is noteworthy that binding of the Pax7 mutants to the group of Resistant sites, binding sites that show no chromatin change upon Pax7 binding, is markedly impaired by the PD mutations but relatively unaffected by the HD S246E mutation. This data indicates that the PD domain is primarily involved in scanning heterochromatin sites for binding and that this function is relatively independent of the HD.

The loss of recruitment observed in ChIPseq (assessed as the presence of peaks above threshold) is significant for the Pioneered, Primed and Activated sites but it is not complete (Fig. 4B). In order to ascertain the meaning of this residual recruitment, we selected subsets of enhancers (Constitutive, Pioneered or Activated, Fig. 4 D-F) based on presence of a single Pax7 recognition motif, either composite, PD or HD motif. For comparisons, average profiles of Pax7 ChIPseq peaks containing multiple motifs are shown in Fig. 4C. The Constitutive Pax7 peaks are relatively insensitive to either PD or HD domain mutations irrespective of the presence of multiple motifs (Fig. 4C) or of only one motif (Fig. 4D-F). In order to correlate Pax7 recruitment with pioneering activity, we performed ATACseq for the Pax7 DBD mutants and compared them to the ATAC profiles of cells expressing wild-type Pax7. These profiles (Fig. 4C-F, bottom panels) illustrate the open chromatin status of Constitutive enhancers independently of Pax7 recruitment.

In sharp contrast, the Pioneered peaks show dependence on intact DBDs for both PD and HD, as much for Pax7 recruitment as for chromatin opening (ATAC).

Recruitment to the Activated subset is more dependent on DNA recognition specificity. Indeed, peaks that have either single composite (Fig. 4D) or PD motifs (Fig. 4E) show relatively unaltered Pax7 recruitment for the HD mutant S264E but this is only marginally reflected in ATACseq signals which are similarly decreased by both PD and HD mutations. For sites that have a single HD motif, both Pax7 recruitment and ATAC signals are affected by mutations in either PD or HD. Specific loci examples of these effects are provided in Fig. S4. Collectively, these results indicate that PD and HD domains are required to stably bind Pioneer sites and trigger chromatin opening. For Activated sites that already exhibit some chromatin opening (ATAC signal), there is a similar dependence on both domains for binding targets with a single HD motif whereas targets with single PD or composite motifs show dependence for recruitment only on the intact PD but not an intact HD.

We then asked whether gene activation shows similar dependence on intact PD and HD domains as revealed in ChIP recruitment. This was assessed by RT-qPCR measurements of mRNA levels for gene targets that are either dependent on regular transcriptional activation or on pioneering mechanism (Fig. 5A and S5A). Consistent with the Pax7 recruitment and ATAC data, activation/transcription of genes dependent on pioneer action show dependence on both intact PD and intact HD. Activation of gene transcription for gene targets of the Activated class of enhancers is more heterogeneous with some showing dependence on both DBDs (*Pcdh9, Pde2a, Gas6*) and some with lesser dependence on the HD domain (*Gprc5, Pgc*). Whereas virtually all Pax7 target genes are regulated by Pax7-dependent enhancers, one notable exception is the *Oacyl* gene where Pax7 exerts its pioneer action at the promoter (Fig. S5B); this gene shows a quicker response to Pax7 expression compared to most other Pax7 pioneered target genes (Fig. S5A).

**Fig. 5.**
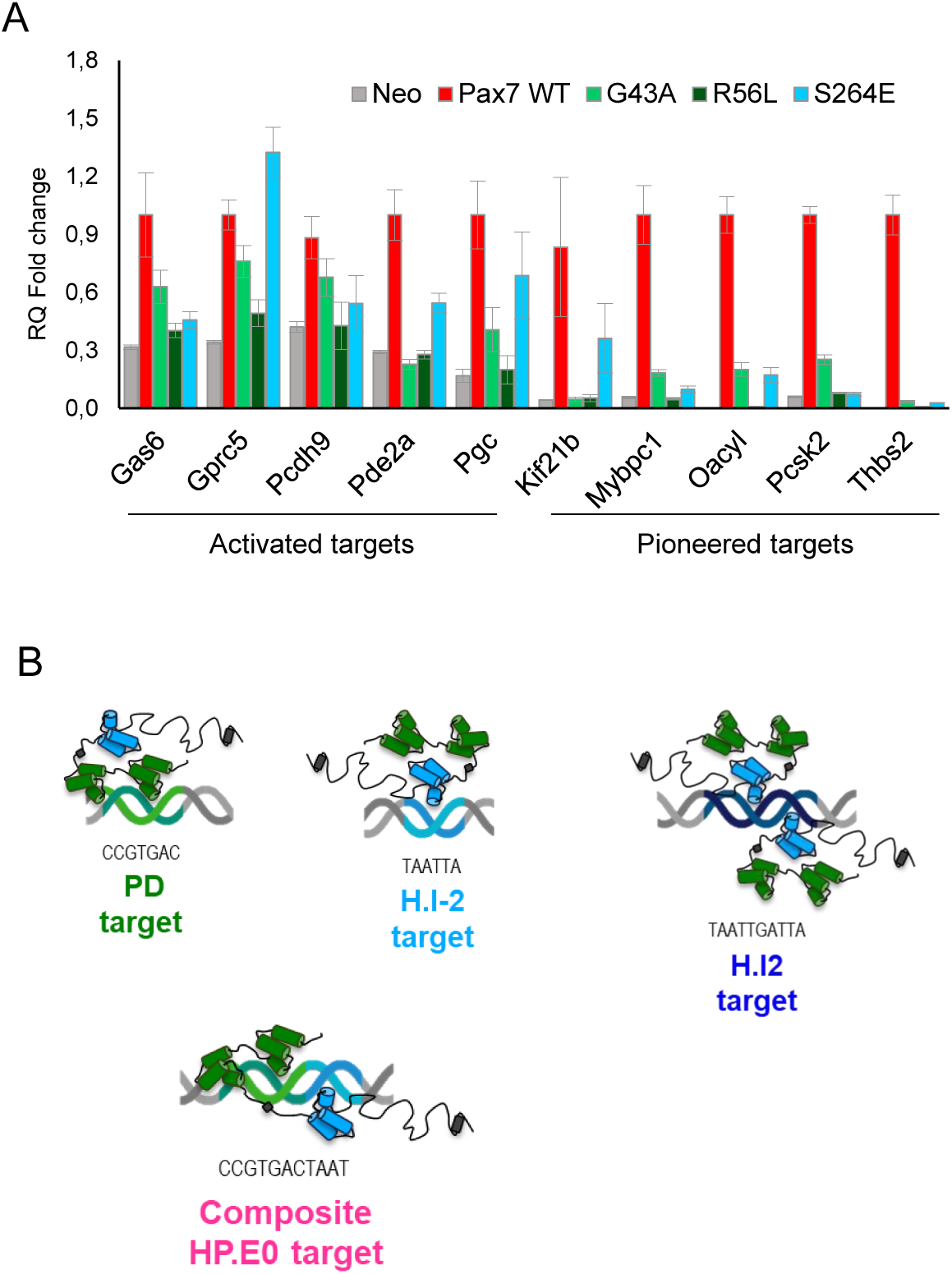
Mutations of either the PD or HD prevent gene activation at targets that require pioneer action whereas they have variable effects on transcriptional target genes. (A)RT-qPCR measurements of mRNA levels for genes subject to activation through either Pax7 pioneering or regular transcriptional activation as indicated. Data are presented for wild-type Pax7 in comparison with mutations in either PD or HD DNA binding domains. (B) Models of Pax7 interaction with its different DNA target sequences.

Collectively, the mutant analyses reveal a strong dependence on both the PD and HD for pioneer activity independently of underlying DNA motifs present in pioneered enhancers. This may relate to a unique arrangement of the PD and HD consistent with interaction at composite binding sites.

## Discussion

This work has queried the requirements for selection of DNA target sites susceptible to pioneer factor chromatin opening. By comparing the DNA sequence motifs present under pioneered sites with those of other subsets of Pax7 binding sites together with mutations within each Pax7 DBD, the unique requirements for pioneer action suggest that structural features required for pioneer activity are far more stringent than for recruitment at enhancers within pre-opened chromatin. Much more than simple DNA sequence recognition is at stake for recruitment of TFs and pioneers, whether considering recruitment of the former at different enhancer states or, of the latter, in different chromatin environments.

What may distinguish the Pax7 pioneered sites compared to other binding/recruitment sites? Compared to all other Pax7 binding sites, the pioneered subset is enriched in high affinity binding sites such as the composite motif, but this site is only present in about 20% of all pioneered enhancers. The pioneered sites are also particularly enriched in the PD motif but this enrichment is not strikingly different when compared to its abundance in other subsets of Pax7 peaks. Nonetheless, Pax7 binding is clearly much stronger at the pioneer subset compared to other subsets (4). Thus, it may be the number of motifs that results in stronger recruitment, for example when comparing the pioneered subset with the constitutive enhancers, but it is not the specific DNA sequence of the motifs present in different subsets that account for their different properties. This argument is strongly supported by the comparison of specific motifs and their abundance in the subset of Resistant compared to Pioneered sites: both subsets have very similar distribution of all PD and HD binding motifs. This observation clearly points towards heterochromatin organization as the defining property that establishes susceptible and resistant heterochromatin for pioneering.

A unique property of pioneer factors appears to be their ability to gain access to their target DNA sequence within nucleosomal and compacted heterochromatin. The molecular basis for this ability remains unexplained although some hypotheses were put forward. For example, it was proposed that FoxA pioneers may have this ability through molecular mimicry where a portion of the winged helix domain may mimic a structure within linker histone H1 (7). However, this model is not likely to apply to many other pioneers. The Foxa pioneers also have the ability to bind low-affinity sites within heterochromatin and the ability to recognize these low sequence conservation sites was proposed to represent a mechanism to scan putative sites within heterochromatin (21). Further, the presence of a core histone interaction domain within FoxA factors (23) clearly supports the hypothesis of direct chromatin interaction as a unique feature of pioneers providing them access to heterochromatin. Be that as it may, this fails to adequately explain how specific sites are selected for pioneering. It is quite striking that the ability for Pax7 recruitment to the weak binding sites of the Resistant subset is almost entirely dependent on the PD, but not the HD (Fig. 4B). This preference is not reflected in an enrichment for the PD motif within the Resistant subset compared to HD motifs and thus likely reflects a property of the Pax7 PD to recognize heterochromatin.

Whereas the PD appears predominantly responsible for scanning heterochromatin sites, chromatin opening and gene activation by Pax7 pioneering is completely dependent on both intact PD and HD. This dependence is clear for both the fully activated pioneered sites as well as for the primed sites: hence, this dependence probably reflects requirements for the initial step of the chromatin opening mechanism. The dependence on both intact DBDs for pioneering is not strictly correlated with the presence of cognate DNA sequence motifs for these domains within the pioneered subset of enhancers, although this subset has the highest enrichment for all motifs and particularly the high-affinity composite (HP.E0) sites. Given the higher affinity of the composite site for Pax7 binding in vitro and its association with a Pax7 monomer (Fig. 5B), but recognizing that many pioneered enhancers do not contain this high-affinity site, it is reasonable to propose that a unique conformation of the two DBDs relative to each other is associated with pioneering ability, and that this conformation can be achieved thought different combinations of DNA sequence motifs arrangement. This hypothetical conformation illustrated over the composite motif may differ compared to the Pax7 dimers interacting with the palindromic H.I2 target or even compared to simple PD or HD motifs (Fig. 5B).

The situation at pioneered enhancers is in striking contrast with the absence of dependence on either intact PD or HD for recruitment to the Constitutive subset of enhancers that were already in an open and active chromatin state before Pax7 expression. This independence towards intact DBDs is correlated with the lesser abundance of PD and HD motifs within the Constitutive subset of enhancers and likely reflects the greater importance of protein-protein interactions at these enhancers that are already occupied by an array of different transcription factors. It is noteworthy that the subset of Pax7 targets that exhibits the greatest dependence on cognate DNA sequence motifs are found within the Activated subgroup of enhancers, those enhancers that are in a primed state at the onset but become fully activated upon Pax7 binding as revealed by their recruitment of the co-activator p300. For this subset, the presence of a cognate DNA motif is most of the time correlated with the loss of this binding for the Pax7 mutant that contains a substitution in the corresponding DBD. It is noteworthy that target DNA sequence recognition is most important for this group of enhancers that are awaiting for a cognate TF for full activation. In this instance, the pioneer properties are likely not required as the chromatin is already slightly accessible, ie in the primed state.

In summary, the present work defined unique contributions of the two Pax7 DBDs towards pioneer action. First, pioneer factors demarcate themselves compared to other TFs by the ability to recognize targets within heterochromatin and this property appears to be associated with the PD. Second, pioneered sites exhibit the strongest recruitment in ChIPseq and this correlates with higher frequency of cognate DNA motifs and of sites of high affinity. Finally, a unique conformation that requires both the PD and the HD of Pax7 is required for chromatin opening at pioneer sites and gene activation.

## Materials and Methods

### Cell culture

AtT-20/D16v-F2 cells were cultured in Dulbecco’s modified Eagle’s medium supplemented with 10% fetal bovine serum and antibiotics (penicillin/streptomycin). To generate stable transgenic AtT-20 Pax7 G43A, R56L and S264E cell populations, retroviruses were packaged using the Platinium-E Retroviral Packaging Cell Line (Cell Biolabs, catalog #RV-101) and infections performed as described (28). Selection of retrovirus-infected cell populations was achieved with 400 μg/mL Geneticin (Gibco, #11811-031). Resistant colonies were pooled to generate populations of hundreds of independent colonies.

### Electrophoretic mobility shift assays (EMSA)

Gel shift assays were performed using 1 μg of nuclear protein extracts from different AtT-20 cell lines pre-incubated 10 min on ice with 1 μg of nonspecific carrier DNA poly(dI-dC) and poly(dA-dT) in the binding buffer (25 mM HEPES pH 7.9, 84 mM KCl, 10% glycerol, 5 mM DTT). If applicable, cold competitor probe was added to the pre-incubation binding mix. Equal amounts of radiolabeled DNA probe (50,000 cpm) were then added for a total volume of 20μL and incubated 60 min on ice. If applicable, 1 μg of Flag M2 antibody (F3165, SIGMA) was added to the mix during the last 10 min of the incubation.

#### Electrophoresis

Nondenaturing polyacrylamide gels (40 mM Tris, 195 mM glycine, 0.08% APS, 0.5 μL/mL TEMED, 5% Acrylamide:Bisacrylamide in 19:1 ratio), were prerun in Tris-glycine buffer (40mM Tris and 195 mM glycine) for 60 min at 300 V at 4°C. The gel shift reaction mixture was then loaded, and the gel was run for 3 h at 300 V at 4°C. Gels were then dryed on 3MM Whatman paper in a gel dryer at 80°C during 60 min under vacuum. Autoradiography films (HyBlot CL, catalog # E3018) were then exposed with the dried gels for 10 to 48 h using an intensifier screen.

### ChIPseq and ATACseq

ChIP experiments were performed in AtT-20 cells as previously described (29) using 5 μg of antibody (anti-FLAG-M2, mouse monoclonal, Sigma F3165). ATACseq was performed as previously described (4). Relative Pax7 ChIP and ATACseq read density profiles were generated by Easeq average signal function (30) (http://easeq.net) and data were exported to Microsoft Excel to create graphs. ChIPseq and ATACseq heatmaps were generated by Easeq. To visualize ChIPseq and ATACseq profiles, we used the Integrative Genome Viewer tool (31).

Supplementary Information on Materials and Methods are available.

## Acknowledgements

We are grateful to Odile Neyret and her group at the IRCM Sequencing Core Facility for NGS analyses and to Valerie Magoon for her expert secretarial assistance. Data analyses were possible thanks to the support of Compute Canada. This work was supported by a Foundation grant (FDN154297) to J.D. from the Canadian Institutes of Health Research.

The authors declare no conflict of interest.

## Supplementary Information

### Supplementary Information on Materials and Methods

#### Expression and purification of Pax7 polypeptides from bacteria

For the expression of polypeptide fragments of Pax7 in bacteria, the sequences encoding the PD alone (PD; residues 35-158 of mouse Pax7 Uniprot P47239-1),the HD alone (HD; residues 215-273 of mouse Pax7), the PD plus the linker region (PD-L; residues 35-215 of mouse Pax7) or the PD plus linker plus HD (PD-HD; residues 35-273 of mouse Pax7) were incorporated into the pET15b (Novagen) vector. The vectors were then transformed in *E. coli* host strain BL21DE3 (Novagen) and the cells were grown at 37°C in Luria Broth (LB) media overnight, diluted 1:4 with fresh LB and expression of the His-tagged Pax-7 polypeptides was induced for 4h at 30°C with 0.7 mM isopropyl-β-D-thiogalactopyranoside (IPTG; Inalco). The cells were harvested by centrifugation, suspended in lysis buffer (either 20 mM Tris-HCl pH 7.4, 1M NaCl and 50 mM imidazole for HD, PD-L and PD-HD or 20 mM NaPO4 pH 7.4, 1M NaCl and 50 mM imidazole for PD), lysed with a French press and centrifuged at 105,000g for 1h at 4°C. The supernatant was collected and incubated with a nickel charged Chelating-Sepharose resin (GE healthcare) for 1h at 4°C. Following incubation, the resin was thoroughly washed with 20 mM Tris-HCl pH 7.4, 1M NaCl and 50 mM imidazole, and the polypeptides wer eluted from the resin with elution buffer (20 mM Tris-HCl pH 7.4, 1M NaCl and 500 mM imidazole). The eluted proteins were dialyzed against 20 mM Tris-HCl pH 7.4, 1mM EDTA buffer overnight and further purified on a 50 mL High Performance SP-Sepharose (GE Healthcare) column using a gradient with 20 mM Tris-HCl pH 7.4, 0.75 M NaCl,1mM EDTA over 5 column volumes. The fractions containing the purified polypeptides were dialyzed against 20 mM Tris-HCl pH 7.4, 1mM EDTA buffer overnight and stored at -80 °C until usage.

#### Electrophoretic mobility shift assays (EMSA)

##### Probe labeling

Double-stranded DNA probes were prepared by annealing of oligonucleotides (1 μM of each: see sequences in the Supplementary Figure 1A) in buffer TEN (10 mM Tris pH 8.0, 1 mM EDTA and 50 mM NaCl). Labeling was done at room temperature for 20 min in the following buffer: 50 mM Tris pH 7,5, 10 mM MgCl2, 1 mM DTT and 0.5 mg/mL BSA supplemented with 0.25 μM dGTP, dTTP, 0,2 μM α-^32^P-dCTP (3,000 Ci/mmol, 10 mCi/mL, Perkin Elmer) and 1 u of DNA polymerase I Large (Klenow) fragment from *E. coli* (NEB, catalog # M0210M). Probes were then purified on Micro Bio-Spin P-30 Tris chromatography column (Bio-rad, catalog # 732-6223).

##### Nuclear extracts

To prepare nuclear extracts, AtT-20 cells from a 100mm petri dish were washed and collected in PBS. Pelleted cells were resuspended in 800μL of cold low-salt buffer (10 mM HEPES pH 7.9, 10 mM KCl, 0.1 mM EDTA, 0.1 mM EGTA, 1 mM DTT, 0.5 mM PMSF, 0.6 μM aprotinin, 0.6 μM leupeptin and 2 μM pepstatin A). After 15 min incubation on ice, 100 μL NP40 10 % was added. Cytosolic fractions were discarded, and nuclear proteins extracted in 100 μL of cold high-salt buffer containing 20 mM HEPES pH 7.9, 400 mM KCl, 1 mM EDTA, 1 mM EGTA, 1 mM DTT, 1 mM PMSF, 0.6 μM aprotinin, 0.6 μM leupeptin and 2 μM pepstatin A. Protein concentration was quantitated by Bradford assay.

### NGS data analysis

ChIP-seq and ATAC-seq paired-end sequenced reads were trimmed using Trimmomatic/0.36 (1) and aligned to the mouse mm10 reference genome using Bowtie/2.3.5 (-- very-sensitive --no-mixed --no-unal) (2). Bam files were created using view from SAMtools/1.9 (3) and duplicated reads were removed using MarkDuplicates from Picard/2.17.3 (Broad Institute). Coverage tracks were created using bamCoverage (--normalizeUsing RPKM -bs 10 - e) from deepTools/3.3.0 (4). Peaks were called against sequenced input DNA or control Flag ChIP using callpeaks (-f BAMPE -p (1e-3) -g mm) from MACS/2.1.1 (5). Categories of Pax7 binding sites are based on already published lists (6) but refined as described in Supplementary Table 1 to increase their homogeneity. Pax7 ChIP-seq peaks were categorized using the absence or presence, before or after Pax7 expression, of overlapping peaks from: H3K4me1 ChIP-seq to assess enhancer marking; p300 ChIP-seq to assess enhancer activation; and ATAC-seq to assess DNA accessibility. SMC1 ChIP-seq was used to avoid insulator contamination in the resistant category. In order to minimize threshold effects, the absence of a peak requires no peak to be detected with a low detection threshold whereas its presence requires its detection with a higher threshold.

### Motifs analysis

*De novo* motif analysis of Pax7 ChIP-seq data was performed on 60 bp surrounding peak summits. Each subset of Pax7 peak were processed using HOMER *de novo* motif search (7) and MEME (http://meme-suite.org/tools/meme-chip), in order to find the motifs with the most significant P-value. Graphical representations of the position weight matrices obtained from these analyses were generated using a custom Python script for motif logos. Motifs frequencies were analyzed using a custom Python based program. Using uncovered matrices, motif scores for each Pax7 binding site were computed with a custom Python script in a 100 bp window around Pax7 peak summits.

### Whole-genome bisulfite sequencing (WBGS)

The CpG methylation levels were represented using the published data from (6).

### RT-qPCR

RNA was extracted from AtT-20 cells using the RNeasy Mini kit (Qiagen #74104), and cDNA as synthesized using SuperScript III reverse transcriptase (Invitrogen, #18080-044) following manufacturer’s recommendations. The resulting cDNAs (5 μl of 1/50 dilution) were analyzed by qPCR using SYBR Green reagent (ThermoFisher Scientific #A25741) supplemented with 500 nM of each gene-specific primer pair (provided in Supplementary Table 2) in a total volume of 10 μl. At least 3 biological replicates were analyzed in duplicates for each condition using a ViiA™ 7 Real-Time PCR device (ThermoFisher scientific), and results were analyzed using the accompanying software. Gene expression values were normalized to those of GAPDH and the statistical significance assessed using bilateral Student’s T-test with unequal variances on Microsoft Excel.

## Legends to Supplementary Figures

**Supplementary Figure 1.**
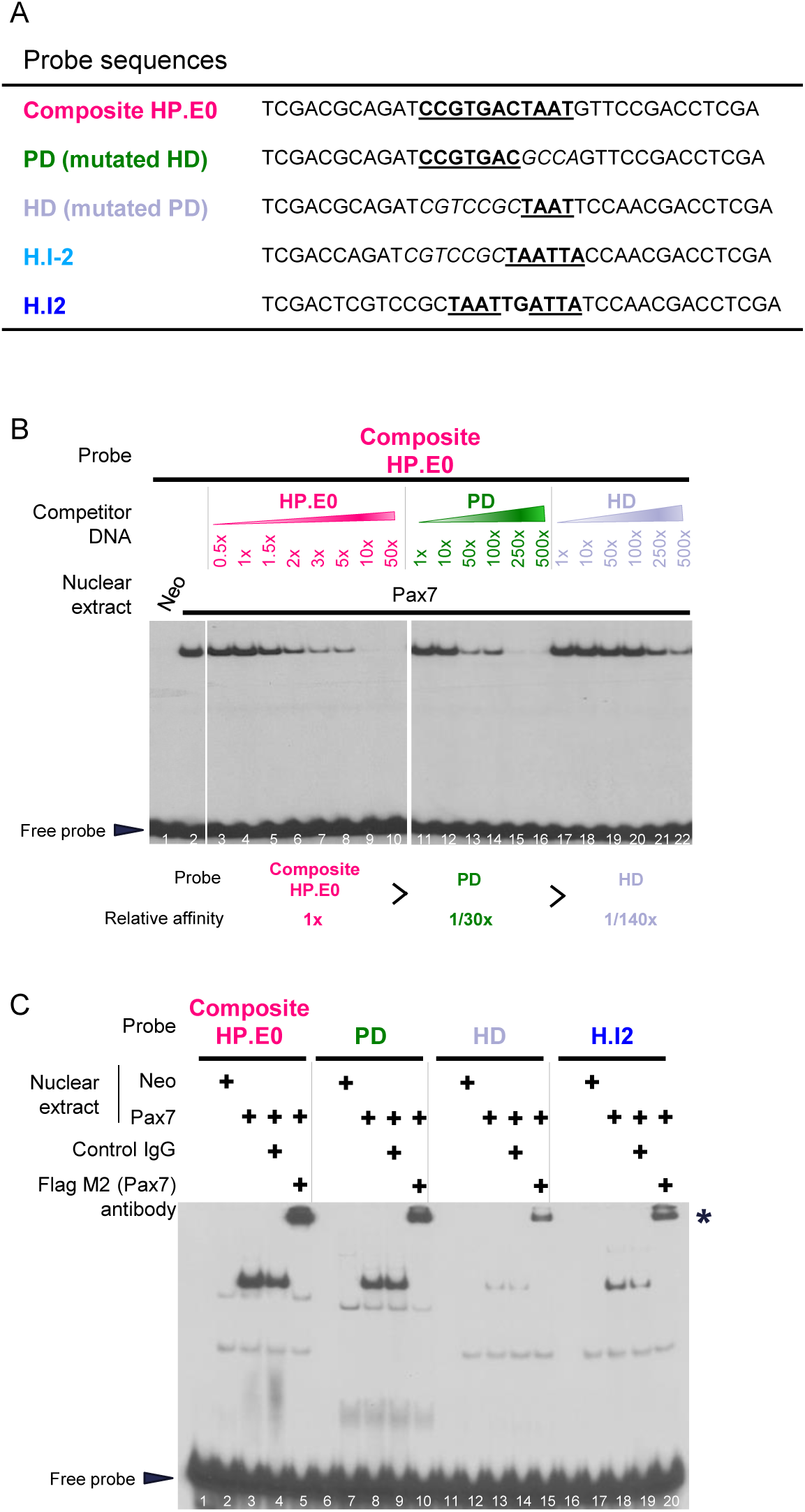
Characterization of Pax7 EMSA complexes with various probes. A) DNA sequences of probes used in gel retardation assays (EMSA). B) Competitive EMSA assays to assess relative affinities of composite HP.E0, PD and H.I2 targets. Complexes formed between the HP.E0 probe and Pax7 from AtT20Pax7 nuclear extracts were competed with the indicated molar excesses of either HP.E0, PD or H.I2 probes. C) EMSA supershift experiments using the Flag2 antibody to supershift Flag-Pax7-containing DNA complexes.

**Supplementary Figure 2.**
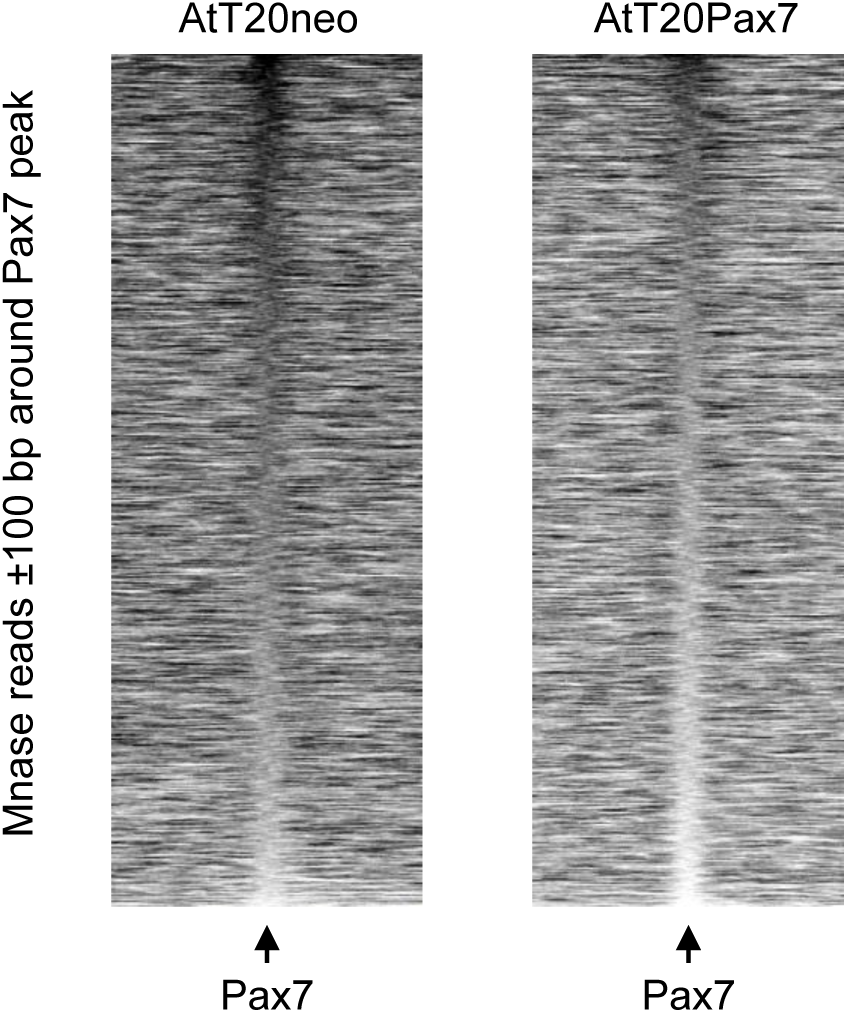
Nucleosome positioning relative to Pax7 Pioneer sites was determined by MnaseSeq analyses in AtT20neo and AtT20Pax7 cells. The heatmaps represent read densities +/- 100 bp on either side (up to +/- 1000bp) of the sites pioneered by Pax7 in AtT20 cells.

**Supplementary Figure 3.**
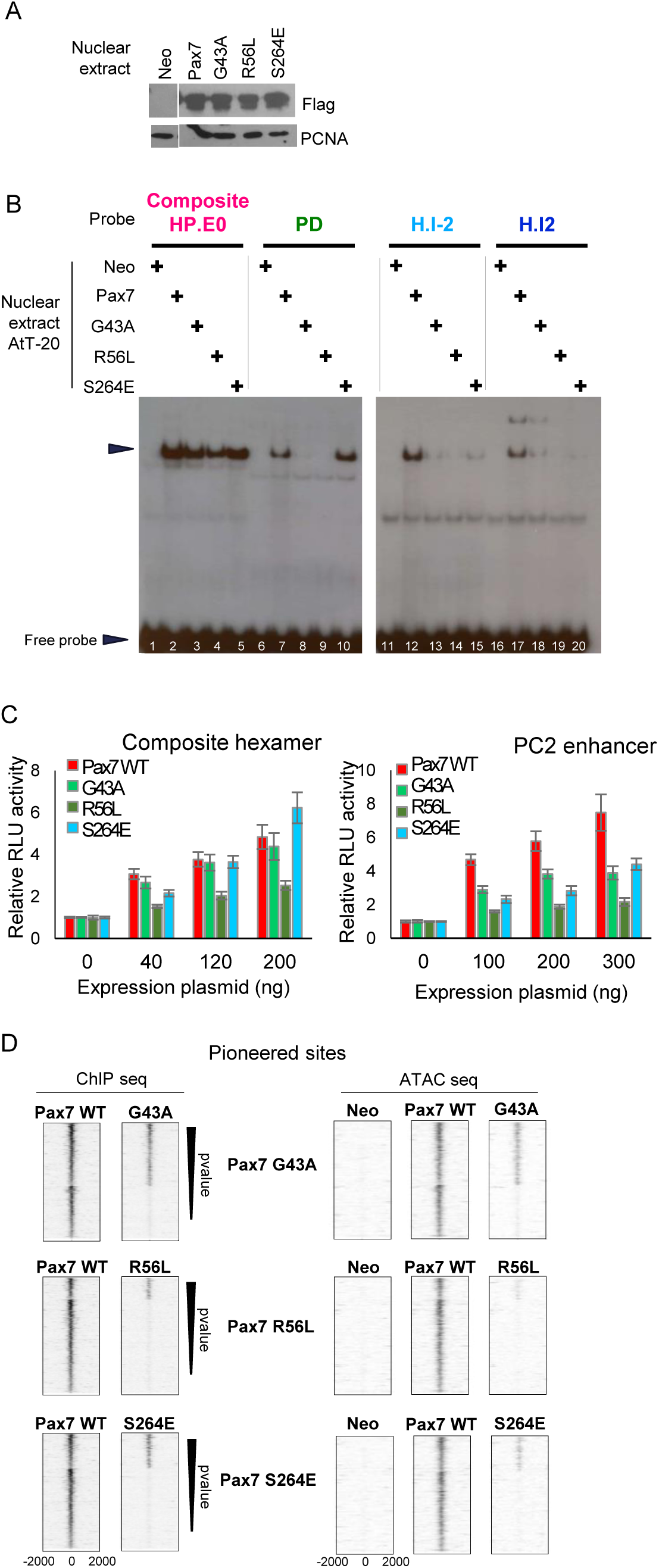
Characterization of Pax7 mutants within the PD (G43A and R56L) and HD (S264E) domains. A) Western blot analysis for expression of Pax7 and three mutants in AtT20 cells expressing N-terminally flagged Pax7 proteins. PCNA Western blot is used as internal control. B) EMSA using the indicated DNA probes incubated with nuclear extracts from AtT20 cells expressing the indicated Pax7 and mutants. C) Transcriptional activity of Pax7 and its DBD mutants assessed by cotransfection of indicated expression plasmids together with a luciferase reporter plasmid driven by six copies of the HP.E0 composite or by the PC2 gene enhancer. Luciferase activity was assessed as previously (4) in extracts of AtT20 cells prepared 36 hours post-transfection with Lipofectamine. D) Genomic recruitment (Flag-Pax7 ChIPseq) and ATACseq profiles at the Pax7 Pioneered sites compared to the same sites in AtT20 cells expressing the three Pax7 DBD mutants.

**Supplementary Figure 4.**
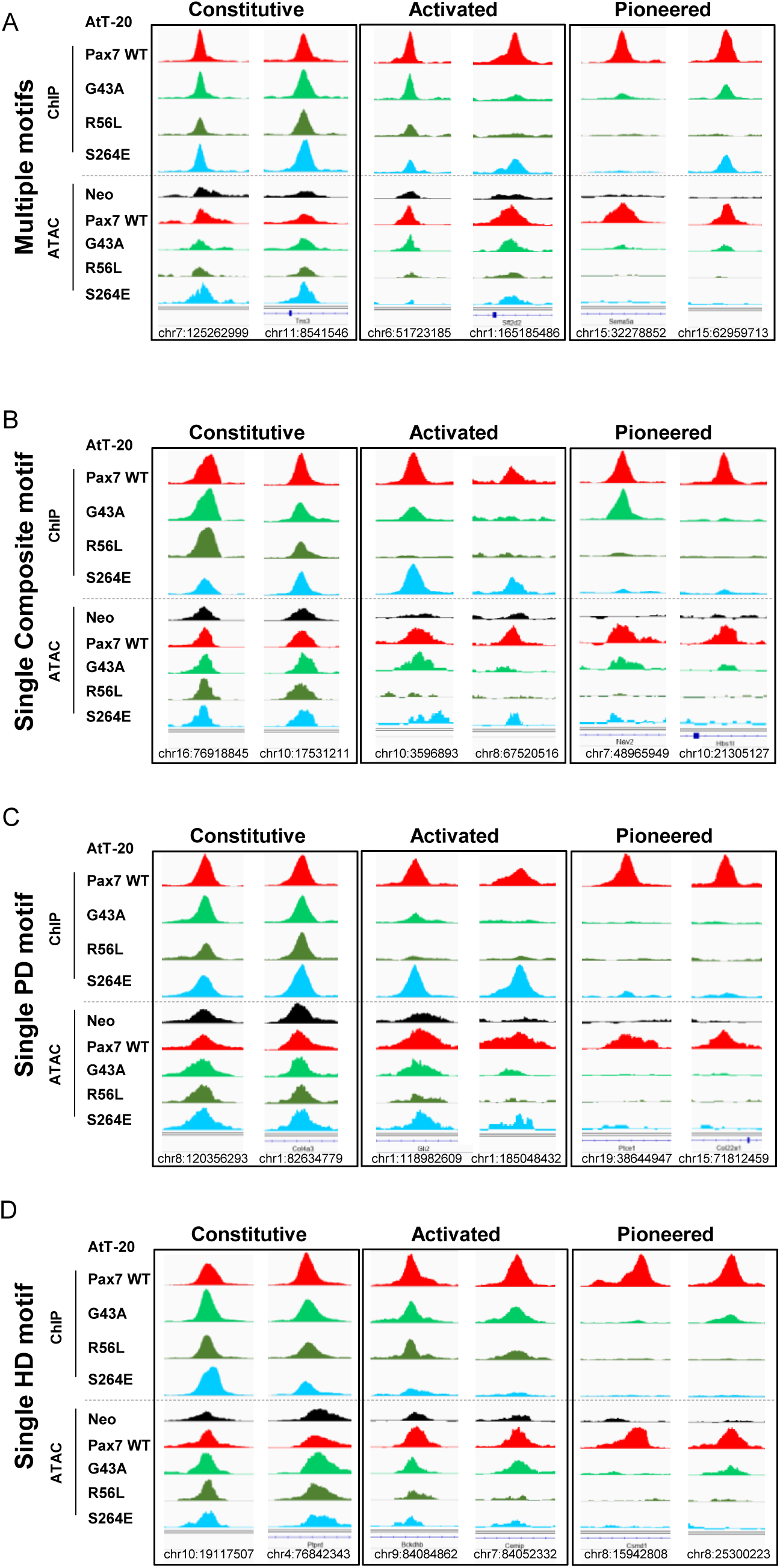
Representative loci for genomic recruitment (Flag-Pax7 ChIPseq) and ATACseq for wildtype Pax7 and three DBD mutants as indicated. Two representative loci are shown for each subset of peaks in the Constitutive, Activated and Pioneered categories. Examples are shown for peaks containing multiple motifs (A), or only one Composite (B), PD (C) or HD (D) motif.

**Supplementary Figure 5.**
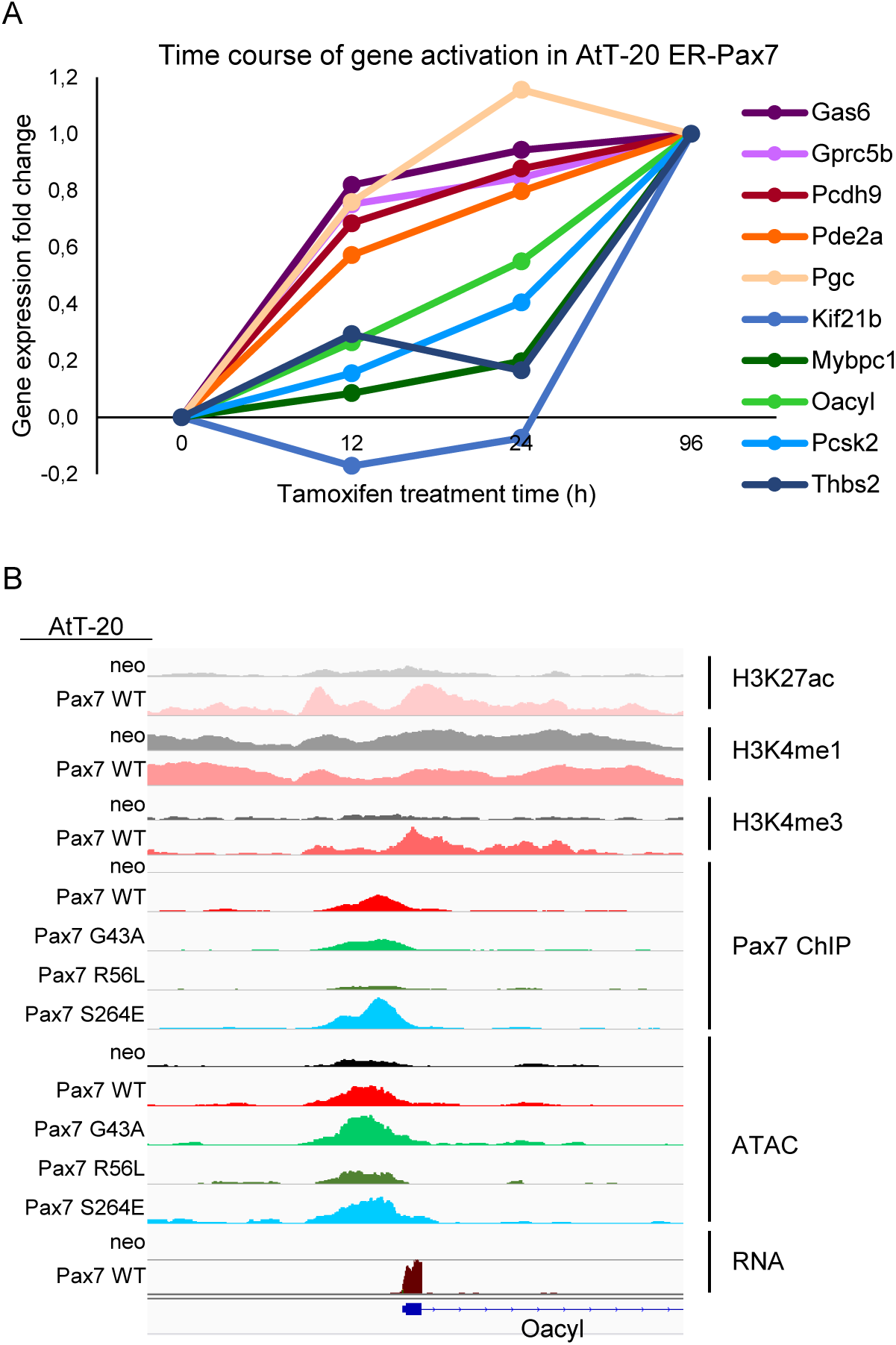
Pax7 target genes. A) Time courses of mRNA accumulation determined by RNAseq analyses (5) showing rapid responses for genes regulated through “Pax7 Activated” enhancers compared to the slower responses of “Pioneered” target genes. B) A unique example, the *Oacyl* gene, of Pax7 pioneer action at promoter rather than enhancer sequences. Gene browser views of the *Oacyl* locus show the indicated ChIPseq and ATACseq analyses performed in AtT20Neo, AtT20Pax7 WT or mutant cells.

**Supplementary Table 1.**
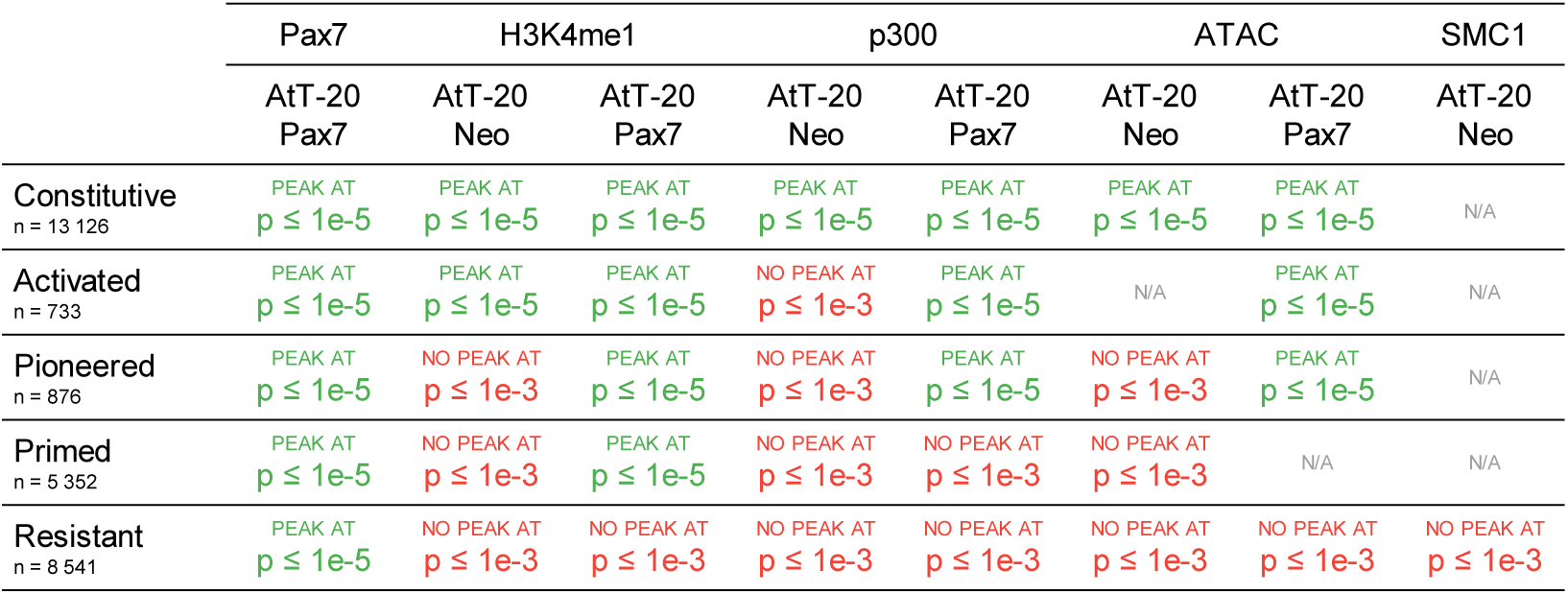
ChIP and ATAC-seq thresholds to categorize Pax7 peaks

**Supplementary Table 2.**
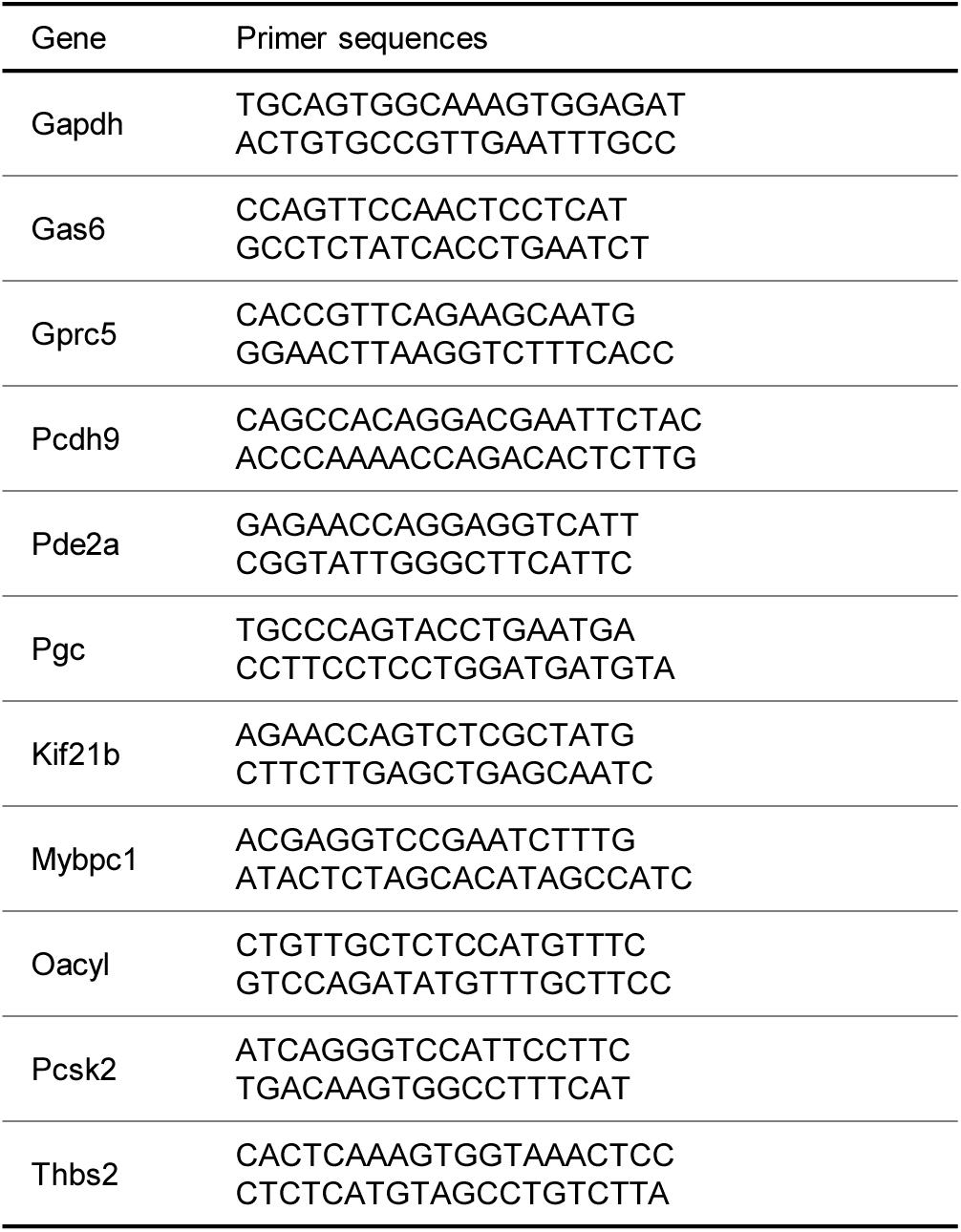
Sequences of oligonucleotide primers

## Notes

### Competing Interest Statement

The authors have declared no competing interest.

